# A computational model of fractal interface formation in bacterial biofilms

**DOI:** 10.1101/2022.05.10.491419

**Authors:** Caelan Brooks, Meiyi Yao, Jake T. McCool, Alan Gillman, Gürol M. Süel, Andrew Mugler, Joseph W. Larkin

**Affiliations:** Department of Physical Sciences, Kutztown University of Pennsylvania, Kutztown, Pennsylvania, USA; Department of Physics, Boston University, Boston, Massachusetts, USA; Biological Design Center, Boston University, Boston, Massachusetts, USA; Department of Physics, Harvard University, Cambridge, Massachusetts, USA; Department of Physics & Astronomy, University of Pittsburgh, Pittsburgh, Pennsylvania, USA; Division of Biological Sciences, University of California San Diego, La Jolla, California, USA; Evident Scientific, Waltham, Massachusetts, USA; Department of Biology, Boston University, Boston, Massachusetts, USA

**Keywords:** agent-based, pattern formation, biofilm

## Abstract

Bacteria benefit from cellular heterogeneity: cells differentiate into diverse gene expression states. As colonies grow, cellular phenotypes arrange into spatial patterns. To uncover the functional role of these emergent patterns, we must understand how they arise from cellular growth and mechanical interactions. Here we present a simple, agent-based model to predict patterns of motile and extracellular matrix-producing cells in biofilms of *Bacillus subtilis*. By incorporating phenotypic inheritance, mechanical interactions, and peripheral motile cell dispersal, our model predicts the emergence of a pattern: matrix cells surround a fractal-like interior motile population. We find that, while some properties of the motile-matrix interface depend on initial conditions, the motile distribution at large radii depends solely on the model’s growth mechanism. The phenotypic interface exhibits a fractal dimension that increases as biofilms grow but reaches a maximum as the peripheral layer of matrix cells exceeds the capacity of the inner cells to push it out of the way. By varying parameters, we find correlations between the interface fractal dimension and expansion of motile cells. We validate findings using experiments on *B. subtilis* biofilms in microfluidics. Our model demonstrates the emergence of colony-level phenotypes from single cell-level interactions and cells modifying their own environment.

## INTRODUCTION

In stressful environments, bacteria differentiate into heterogeneous states of physiology and gene expression(1; 2; 3). For example, *Bacillus subtilis* bacteria generate multiple cell types characterised by distinct gene expression states(4), including matrix producers, which synthesise extracellular polymers to bind cells together into biofilms(5) and motile cells, which exhibit swimming and swarming behaviour(6). As cells differentiate into these states, they change how they interact with each other and the environment(7; 8). Surface-adhered biofilm colonies contain cells in numerous phenotypic states(9; 10), leading microbiologists to make analogies between the growth of biofilms and cell differentiation during development of multicellular organisms(11; 12; 13).

Bacterial biofilms are characterized by the synthesis of extracellular matrix, which attaches cells to surfaces and to each other(14). In *B. subtilis*, the matrix-production and motile phenotypes are regulated by a transcription factor network whose interactions cause the two phenotypes to be mutually exclusive(6; 15) and heritable(16). Within *B. subtilis* biofilms, these phenotypic states form repeatable spatiotemporal patterns(10; 17). Despite the robustness of these patterns, we do not understand how they arise. To eventually explore the functional role of spatiotemporal patterns in bacterial colonies, we must develop models that predict patterns based on biological and physical factors that may create them, such as phenotypic inheritance in dividing cells(1) and mechanical interactions among bacteria(18).

Agent-based models provide a framework to ask how colony-scale phenomena emerge from the properties of individual bacterial cells. Such simulations have been used to show how mechanical interactions lead to cellular re-ordering in dense colonies(19), how cell shape affects the spatial partitioning of different species within microbial communities(20), and how mechanical interactions and cell shape together lead to fractal patterning in bacterial layers(21). Furthermore, agent-based simulations have been used to account for feedback of mechanical interactions on cell growth(22) and interplay of mechanics with with other forms of interaction like quorum sensing(23). Nevertheless, the combined effects of mechanical interactions and heritable phenotypic differences in those interactions remain poorly understood.

Here, we present a simple, agent-based, computational model of growing populations of *B. subtilis* to ask how mechanical interactions among phenotypes can drive pattern formation. The model takes into account three key factors in the development of motile-matrix patterning. First, cell phenotypes are mutually exclusive and heritable: cells are either motile or matrix-producing (“matix” for short), with daughter cells perfectly inheriting their parent’s phenotype. Second, motile cells exposed to the edge of the colony will “disperse” and swim away. Third, when a cell on the colony interior divides, it must shove other cells out of the way to make space for the new cell. Matrix cells resist shoving more strongly than motile cells.

From our model’s three simple rules for cellular division, motion, and interactions, we can simulate biofilm growth from an initial inoculum of bacteria. The model produces a biofilm with an interior population of motile cells surrounded by matrix cells. The interface between these two populations develops a fractal edge. We characterise the predicted pattern by computing the fractal dimension of this interface. By varying our model parameters and extracting the fractal dimension of simulated biofilms, we find that our model predicts a correlation between the fractal dimension and the radial expansion of motile cells across a wide range of parameter values. We find that both the fractal dimension and the extent of radial expansion are consistent with experimental data from a *B. subtilis* biofilm grown in a microfluidic device(24). Our model shows how population-level phenomena can emerge from the heterogeneous mechanical cellular interactions.

## RESULTS

### An agent-based, mechanistic cell growth model

We model the dynamics of biofilm growth using sites on a two-dimensional square lattice (Fig. 1). For computational simplicity, we restrict ourselves to parameters for which simulated phenotypic patterns develop when biofilms grow to thousands of sites, whereas experimental biofilms may typically grow to hundreds of thousands of cells. Therefore, each site can be thought of as a coarse-grained packet of cells of the same phenotype (motile or matrix). Furthermore, as described below, we assume no preferential direction for site growth apart from steric hindrance due to existing sites (although we later checked for the effects of preferential growth direction). This assumption avoids lattice artifacts that can arise in agent-based models(25) and is consistent with sites being packets of cells, given that the packets are larger than the length scale over which cells align due to their rod-like shape(26). We later accounted for cellular coarse-graining when comparing to the experimental data. From here on in the paper, we use ‘sites’ and ‘cells’ interchangeably, both referring to a lattice site within the grid.

**Figure 1.**
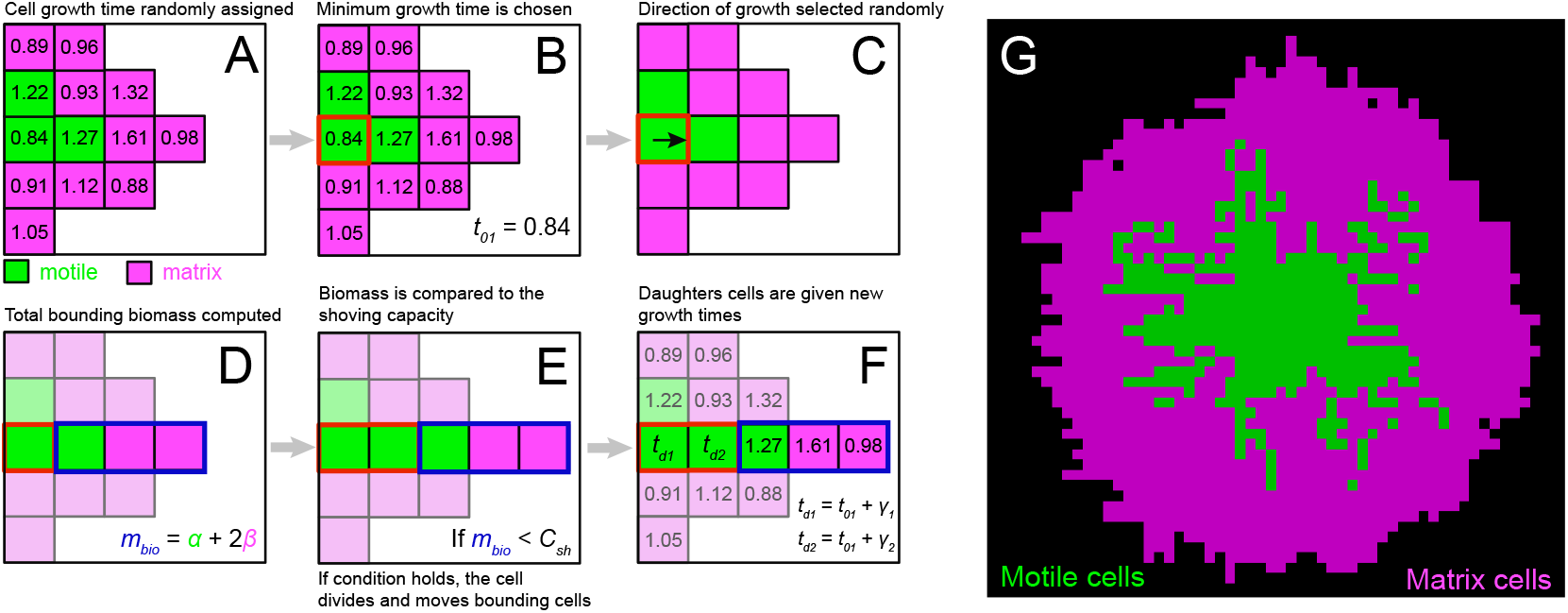
Agent-based biofilm growth model. (A-F) Growth and displacement mechanism: (A) Growth times are drawn from a Gaussian distribution and are assigned to each site. (B) The lattice site with the minimum growth time (outlined in red here) is selected and given the opportunity to grow, where *t*_01_ is the initial growth time. (C) The direction of growth is chosen randomly from four options. (D) If the cell is not on the edge, the total biomass (*m*_*bio*_) will be measured in the growth direction. The biomass of each cell type is represented by *α* and *β*, which are Gaussian-distributed with different means (⟨*β*⟩ > ⟨*α*⟩). (E) The biomass is pushed under the condition *m*_*bio*_ *< C*_*sh*_ (F) New sites are given growth times based on *t*_01_. (G) Resulting pattern once interior motile cells become frozen because biomass exceeds shoving capacity. Parameter values are *C*_*sh*_ = 12, ⟨*α*⟩ = 1, ⟨*β*⟩ = 2, and *σ*_*α*_ = *σ*_*β*_ = 0.1.

We initiate the system by creating an inoculum of cells containing both motile and matrix phenotypes. Each cell is given a growth time that acts as an internal clock, indicating when it is time for the cell to divide (Fig. 1A). These growth times are randomly drawn from a normal distribution centred at 1 (in units of the average cell cycle time), with a standard deviation of 0.1 (the typical order of magnitude of variability in cell cycle duration in *B. subtilis*(16)

At each step of the simulation, the cell with the minimum growth time is chosen (Fig. 1B). The chosen cell then goes through a process of determining if and how to divide and produce a daughter cell. If the cell is on the edge of the biofilm, it chooses from the neighbouring empty spaces with equal probability and grows into one of them, filling the space with a daughter cell. Daughter cells inherit their mother’s phenotype with no switching. The two new cells are then assigned random growth times. If instead the dividing cell is surrounded by other cells (Fig. 1C), then it must shove those cells out of the way to create space for a new cell (Fig. 1D). The direction of growth is chosen with an equal probability of being up, down, left, or right. We then compute the total biomass (*m*_*bio*_) between the chosen cell and the edge of the growing colony in the chosen direction (Fig. 1D). Each cell has a different biomass, with motile and matrix biomasses being drawn randomly from distributions with different means, but the same variance. Motile cell biomass is *α* ±*σ*_*α*_ and matrix cell biomass is *β* ±*σ*_*β*_, where *β* > *α* and *σ*_*β*_ = *σ*_*α*_. This key aspect of the model, having matrix and motile cells contribute differently to mechanical interactions, implements similar concepts to continuum models of phenotypic interfaces, which have shown that purely mechanical effects can drive pattern formation in bacterial communities(27). We hypothesised that different mechanical interactions of motile and matrix cells in our agent-based model would drive the formation of distinct patterns of motile and matrix cells. Because matrix cells have greater biomass in our model, they offer more resistance to pushing by growing cells. This choice reflects the fact that matrix cells produce polymers that adhere them to the substrate and to each other(28).

We further assume a constraint on how deep into the biofilm a cell can grow and shove biomass. Previous agent-based models have used a shoving capacity parameter to represent resistance to pushing by growing cells in agent-based simulations, where cells beyond a certain depth in a colony do not grow and push biomass due to nutrient depletion or mechanical constraints(23). We represent the constraint with a parameter called the shoving capacity, *C*_*sh*_. If the bounding biomass is less than or equal to *C*_*sh*_, then the stack of cells is pushed out of the way, and a new daughter cell is created (Fig. 1E). The original growing cell and the new cell then receive new growth times (Fig. 1F) and the simulation progresses by choosing the cell with the next minimum growth time. Our model differs from previous agent-based models(23) in that motile and matrix cells contribute differently to mechanical resistance, resulting in motile cells being shoved around more during biofilm development.

Finally, if at any step of the simulation there is a motile cell at the edge of the biofilm, it “disperses” and is removed from the colony, which is a reasonable assumption based on previous observation of passive escape of motile cells from the edges of biofilms(29).

Simulating biofilm growth from a small group of initial cells, our model produces colonies with a distinctive pattern of motile and matrix cells. Matrix cells surround a fractal-like interior of motile cells. Fig. 1G shows the output of the model for *C*_*sh*_ = 12, ⟨*α*⟩ = 1, ⟨ *(β*⟩ = 2, and *σ*_*α*_ = *σ*_*β*_ = 0.1. As described later, we found that shoving capacities within the range of *C*_*sh*_ = 12 − 18 produced fractal structures most like those observed in experiments. In our simulations, the interior pattern of motile cells eventually becomes frozen in place after the width of exterior matrix cells exceeds the shoving capacity (Fig. S1, Fig. S2).

Two key assumptions of our model drive salient aspects of the motile-matrix pattern: (1) dispersion of peripheral motile cells creates a motile interior surrounded by a matrix exterior, and (2) differential resistance of the two phenotypes moved motile cells during growth, creating branched arms. We wanted to know how the initial arrangement cells, as well as the values of key model parameters, influenced the predicted motile-matrix patterns.

### Dependence of phenotypic patterns on initial conditions

To investigate how our model’s predictions depend on the initial arrangement of motile and matrix cells, we carried out simulations with three different initial conditions (Fig. 2). In the first case, termed “bullseye,” a layer of matrix cells surrounded a group of motile cells in the initial inoculum (Fig. 2A). In the second condition, “mixed,” the initial colony consisted of a mixture of motile and matrix cells, with each initially occupied lattice site having equal probability of being either type (Fig. 2B). The third condition, “concentric,” consisted of alternating rings of each cell type with matrix cells residing on the outermost ring. We show examples of our model’s simulated biofilm growth from these three conditions in Fig. 2A, B, and C.

**Figure 2.**
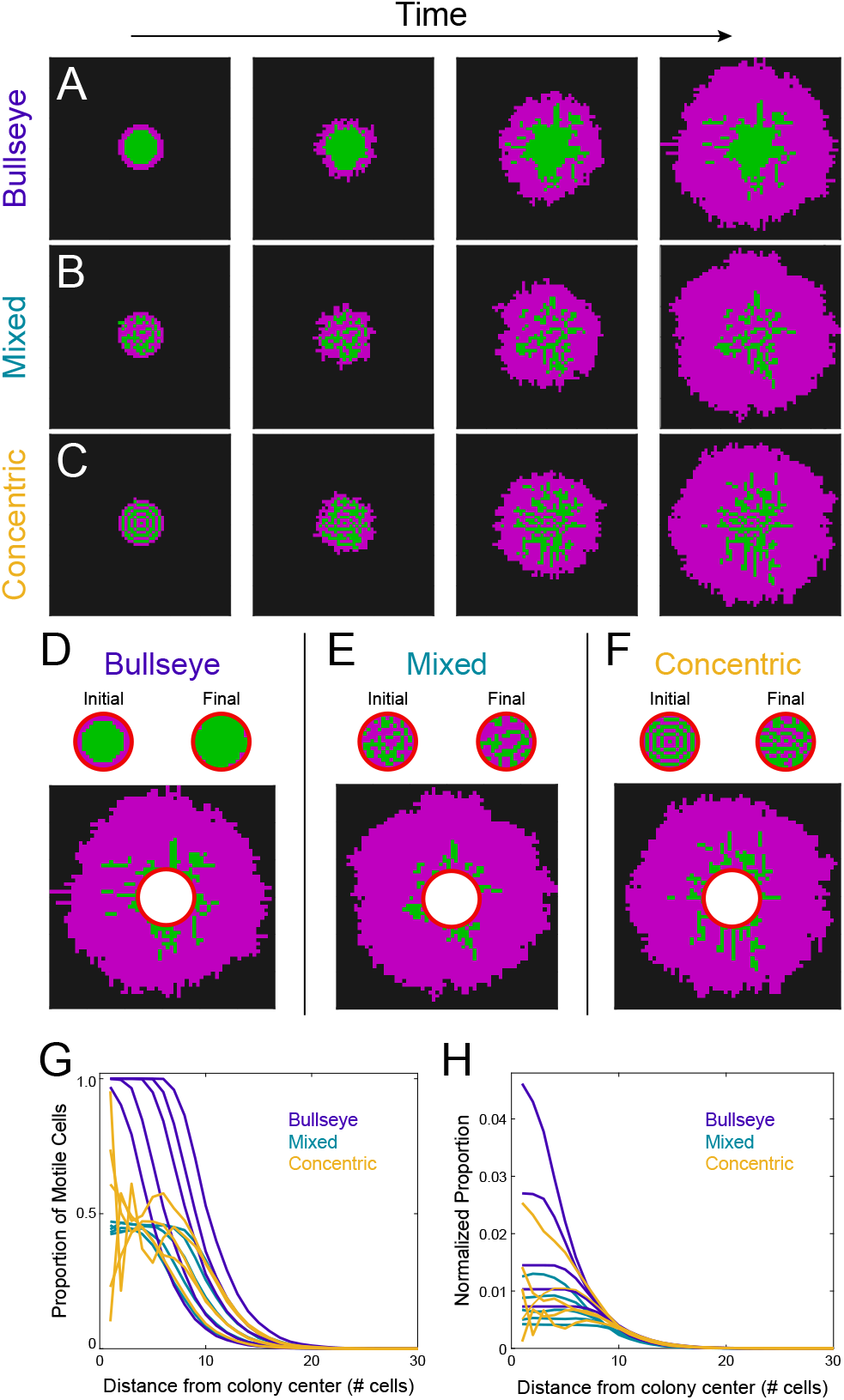
Dependence of patterns on initial conditions. The evolution of the system from three different initial conditions: (A) bullseye, (B) mixed, and (C) concentric. (D-F) Visual isolation of region of the biofilm unaffected by the initial condition. (G) Proportion of motile cells within a ring at a particular distance from the centre of the final pattern, as a function of the distance. Each line within an initial condition subset represents a different initial radius value ranging from *R*_0_ = 4 to *R*_0_ = 8 (larger initial radii correspond to falloffs farther from the centre). (H) When the proportion of motile cells is divided by the initial number of motile cells, a collapse is observed beyond a distance from the centre corresponding to the initial condition. Parameter values are *C*_*sh*_ = 12, ⟨*α*⟩ = 1, ⟨*β*⟩ = 2, and *σ*_*α*_ = *σ*_*β*_ = 0.1.

Each starting condition appeared to create a different final pattern (Fig. 2A, B, and C right). However, we observed that, in the final state for each condition, the regions of the colonies that were populated only by the model’s simulated growth, and not by the initial cells, exhibited qualitatively similar patterns: branches of motile cells that penetrated into an outer layer of matrix-producing cells. We highlight this in Fig. 2D, E, and F. When the initial group of cells is visually excluded, the ability to distinguish which final pattern results from each distinct initial condition is lost.

We next sought to quantify the final patterns of motile and matrix cells generated from each initial condition. We observed that every pattern simulated by our model featured motile cells in the interior surrounded by matrix cells. To quantify these patterns, we computed, for a given ring of cells, the motile cell proportion as a function of the distance of the ring from the colony centre. In Fig. 2G, we plot such curves for bullseye, mixed, and concentric conditions. Each curve is an average of 300 simulations run under the same conditions. Within each initial condition, we analysed three different initial colony radii. The initial proportion of motile cells was kept roughly constant for each initial condition when the inoculation radius changed (with slightly more variation in the bullseye condition than in mixed or concentric). All cases featured patterns where the fraction of motile cells is high in the centre and decays near the edge. However, the curves differed in key ways. For different initial conditions, the fraction of motile cells dropped off at different distances from the colony centre. The same was true for the case of different initial radii within the same initial condition subset: larger initial colonies exhibited a larger radius at which the motile proportion dropped off (Fig. 2G).

Motivated by our observation in Fig. 2D, E, F that biofilm regions that were populated only by model growth looked the same, we thought that perhaps the curves of Fig. 2G would collapse if the initial conditions were taken into account in the proper way. When we normalised the proportion of motile cells throughout different radii by the initial number of motile cells, we produced the curves in Fig. 2H. In this case, we observed a collapse not only of all curves within an initial condition subset, but also of curves representing all initial conditions. The point of collapse of Fig. 2H represents the radius below which the initial condition has an effect. Beyond this point, all normalised curves collapsed, illustrating that our model simulates the distribution of motile cells within biofilms during growth in a way that is insensitive to the initial cell arrangement. This result suggested that the growth dynamics of our model produce patterns of motile and matrix cells that are robust to changes in the initial cell arrangement.

### Fractal interface formation between cell types

In simulated biofilms, the interface between matrix cells and motile cells appeared to have a fractal character (Fig. 1G, Fig 2). Fractal interfaces have been observed in a variety of microbial systems, including the outer edge of bacterial colonies(30) and the trajectories of motile bacteria(31). We chose to quantify the patterns predicted by our model with fractal dimension (Fig. 3A,B). At each time point for model biofilms, we found the outline of the motile-matrix interface and computed its fractal dimension using box-counting(32).

**Figure 3.**
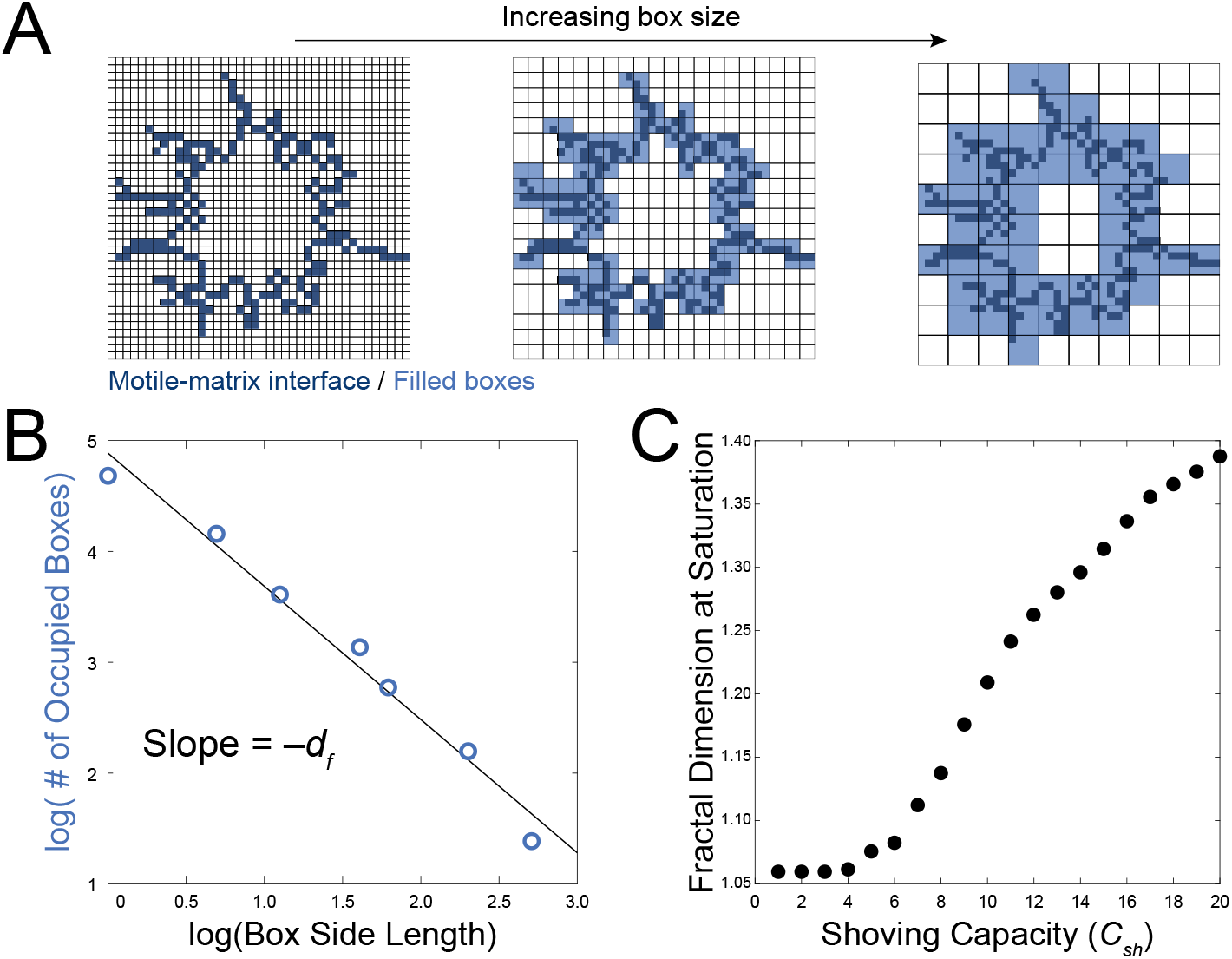
Fractal dimension of phenotype interface. Fractal dimension is found using the box counting method: (A) Grids of different box sizes are fit to the internal motile sub-population to produce a graph like (B). The negative of the slope is the fractal dimension. (C) The fractal dimension at saturation increases with the shoving capacity. Parameter values are, ⟨*α*⟩ = 1, ⟨*β*⟩ = 2, and *σ*_*α*_ = *σ*_*β*_ = 0.1.

In this method, the number of pixels containing a part of the interface is counted. Space is then divided into larger and larger boxes and, for each box size, the number of occupied boxes is counted (Fig. 3A). The number of occupied boxes scales with some power of the box size depending on the shape of the interface. That exponent is the fractal dimension. It is 1 for a simple line, 2 for a two-dimensional plane, and between 1 and 2 for a rough interface like that in our model. To extract the fractal dimension for the interface in our model, we plotted the box size and the number of occupied boxes on a log-log plot and fit a regression line. The negative slope of the regression line is the fractal dimension, *d*_*f*_ (Fig. 3B).

For simulated biofilms initialised with a bullseye pattern, the fractal dimension started out near 1 because the motile-matrix interface was smooth for this condition. Once simulated growth commenced, cells shoved each other out of the way as they grew, roughening the interface, and increasing the fractal dimension. However, once the outer matrix layer thickness exceeded the shoving capacity, *C*_*sh*_, the motile-matrix interface froze into place and the fractal dimension saturated (Fig. S1). Because increasing the shoving capacity increases the time during which cells can grow and push each other around, we hypothesised that the saturating fractal dimension would increase with shoving capacity. After simulating biofilm growth for a range of shoving capacities, we found that saturating fractal dimension does indeed increase with shoving capacity (Fig. 3C). The behaviour of the fractal dimension in Fig. 3C is robust to changes in cell growth decisions, for example whether the direction of cell growth is random or preferential to the previous growth direction chosen by the parent cell (Fig. S2).

### Dependence of fractal dimension on differential biomass of motile and matrix cells

We hypothesised that the phenotypic interface in our model would depend on the difference in biomass between motile and matrix cells. To determine how mechanical differences between motile and matrix cells changed the model’s predicted interface, we simulated colonies with a range of matrix-to-motile average biomass ratios, *ρ*_*βα*_ = ⟨*β*⟩ : ⟨*α*⟩. We found that the interface between cell types became smoother as the matrix-motile biomass ratio increased (Fig. 4A). This makes sense because at very high matrix-motile ratios, interior motile cells can barely push matrix cells at all and the interface remains smooth. In contrast, when both phenotypes have the same biomass (*ρ*_*βα*_ ≈ 1 : 1), motile cells can push matrix cells easily, resulting in a rough interface. To quantify this effect, we computed the interface fractal dimension for a range of *ρ*_*βα*_ values. Specifically, we simulated {⟨*α*⟩, ⟨*β*⟩} = [{1, 1},{1, 2},{1, 3},{1, 5},{1, 10},{1, 100}] with shoving capacities from *C*_*sh*_ = 1 to *C*_*sh*_ = 20. The resulting fractal dimensions are plotted in the heatmap of Fig. 4B. We found that as the matrix-motile biomass ratio increased, the limiting fractal dimension decreased and, above a certain ratio, the interface would not roughen at all. This maximum ratio at which roughening occurs depended on the shoving capacity *C*_*sh*_ (Fig. 4B). Thus, depending on *ρ*_*βα*_ and *C*_*sh*_, the model predicted biofilms that either retained smooth interfaces (dark region of Fig. 4B) or roughened the interface (bright region of Fig. 4B).

**Figure 4.**
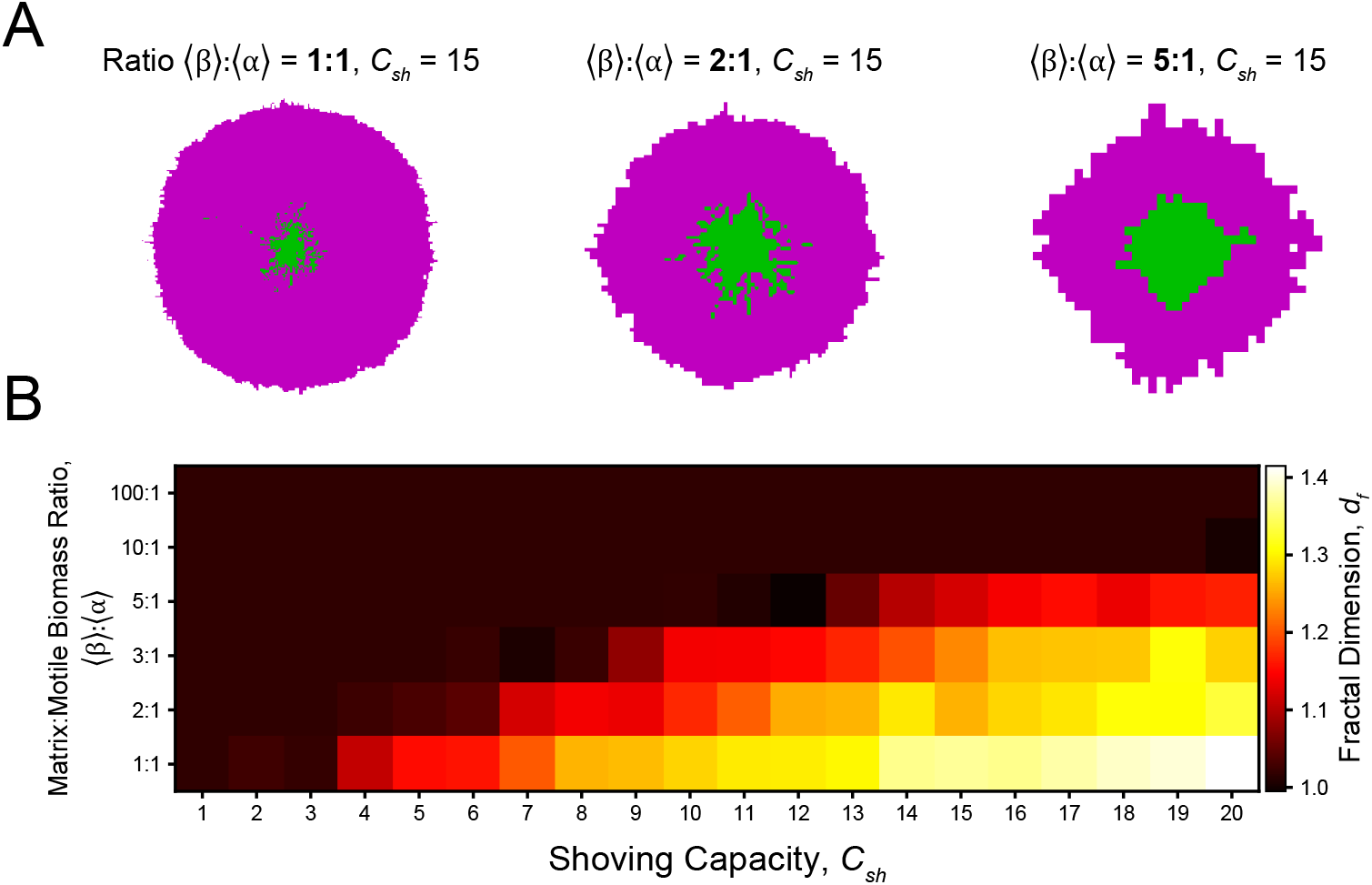
Dependence of fractal interface on motile-matrix biomass ratio. (A) Examples of simulated biofilms with the same *C*_*sh*_, but different ratios of motile to matrix cellular biomass, ⟨*β*⟩ : ⟨*α*⟩. Note that the three examples are depicted at different scales. (B) A heatmap showing fractal dimension for a range of values of *C*_*sh*_ and biomass ratio ⟨*β*⟩ : ⟨*α*⟩. Parameter values are *σ*_*α*_ = *σ*_*β*_ = 0.1.

Our test of parameter dependence (Fig. 4) suggested that the model could predict a wide range of fractal interfaces by simply changing *C*_*sh*_ and *ρ*_*αβ*_. However, we noticed that in simulated biofilms with low fractal dimensions (i.e. smooth interfaces), the fractal dimension froze after relatively little growth (Fig. 4A right). We therefore wondered if there was a relationship between the growth of inner motile cells and the fractal dimension of their interface with matrix cells. To investigate this relationship, we simulated biofilms with a variety of *C*_*sh*_ values, *ρ*_*αβ*_ ratios, and initial cell configurations. We chose initial inoculum conditions that had a range of radii and proportions of each cell phenotype (Fig. 5A). For each simulated biofilm, we extracted the fractal dimension, as well as the average radial expansion of both inner motile cells and the entire biofilm from the start of the simulation to the time when the interface froze (Fig. 5B). After examining fractal dimension and growth for a wide range of model parameters, we found two clear correlations: 1) the radial expansion of motile cells relative to their initial radius (Δ*r/r*, Fig.5B) correlated positively with fractal dimension (Fig. 5C) and 2) the ratio of motile cell radius to total biofilm radius (*r/R*, Fig.5B) correlated negatively with fractal dimension (Fig. 5D). The first correlation arises from the observation that larger Δ*r/r* signifies more growth of motile cells from the initial inoculum. More growth generally means longer fractal motile branches. For the second correlation, a smaller *r/R* ratio is indicative of a larger gap between the internal motile population and the surrounding matrix population, again evidence for longer fractal branches. In contrast, a phenotypic interface closer to the edge of the biofilm results in a larger *r/R*, meaning the motile pattern freezes earlier. In general both correlations fit into the same explanation that if the motile cells at the interface upon initialization have an opportunity to grow, growth events are more likely to occur in these regions as there is less accumulated matrix cells and therefore less biomass to push. The result is motile cells forming branches where longer branches fall into the regime of high Δ*r/r* and low *r/R* where we observe the largest fractal dimensions.

**Figure 5.**
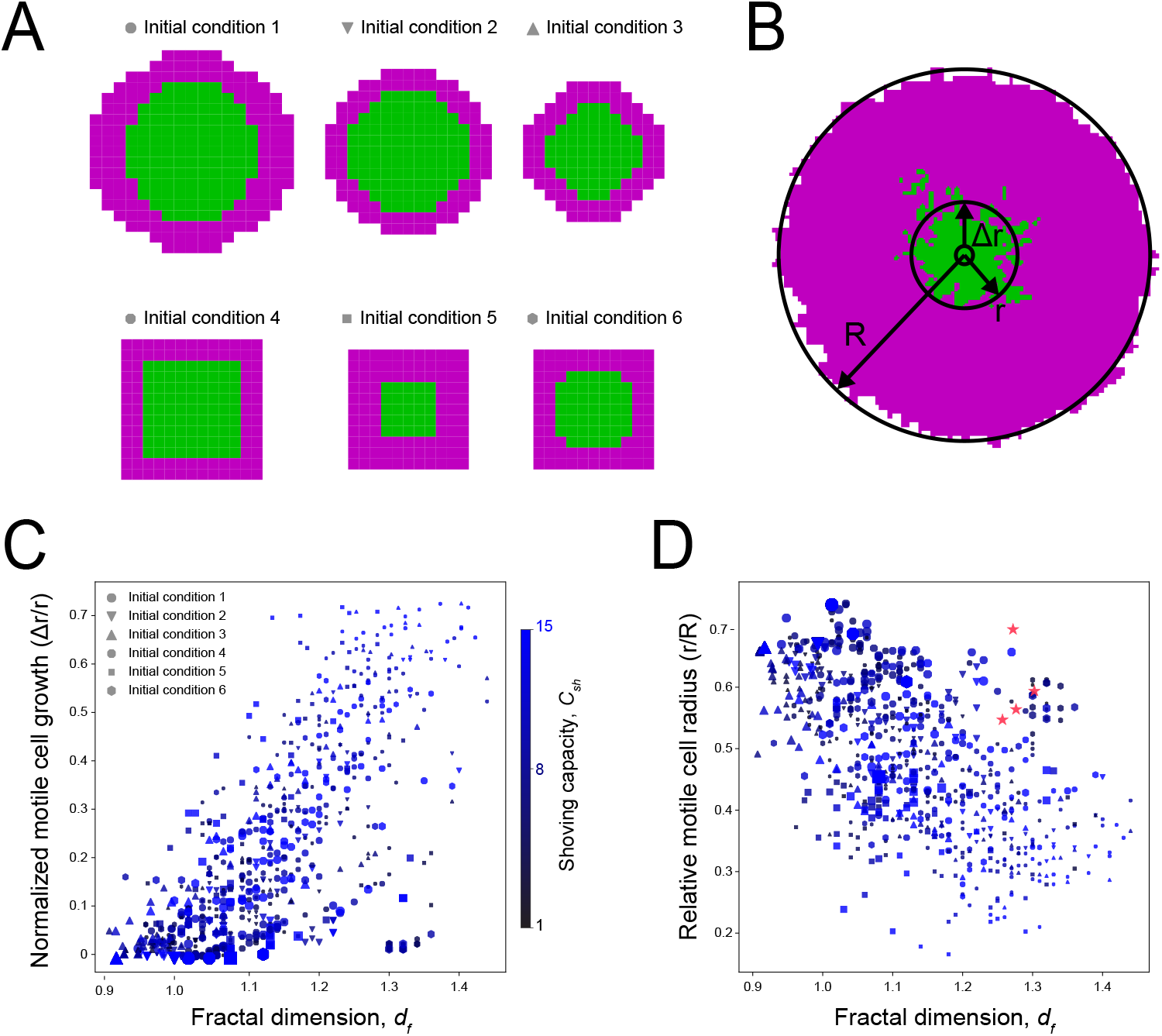
Relationship between fractal interface and biofilm composition. (A) Several different initial conditions simulated for a range of *C*_*sh*_ and biomass ratio values. (B) Schematic shows three parameters of motile and matrix cell radial expansion. (C) Scatterplot of normalized motile cell growth 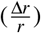 vs. fractal dimension. Marker shape indicates initial cell arrangement, marker size indicates motile-matrix biomass ratio (smallest 1:1, largest 1:10), and marker color indicates shoving capacity. Pearson’s correlation is *r* = 0.70. For each unique set of parameters and initial conditions, 5 simulations were performed, each of which is included in the scatterplot. (D) Scatterplot of the same data for normalized motile cell radius 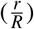vs. fractal dimension. Pearson’s correlation is *r* = −0.65. Parameter values are *σ*_*α*_ = *σ*_*β*_ = 0.1. Pink stars represent experimental data.

These correlations demonstrate general constraints on growth and pattern formation predicted by the model. Specifically, across a broad spectrum of parameters and initial conditions, biofilms that generate fractal interfaces will also exhibit more expansion of motile cells, but those cells will constitute a smaller percentage of the colony once the pattern is formed. Our model predicts a relationship between the mechanical properties of cell phenotypes and the phenotypic content of biofilms formed by those cells.

### Emergence of motile-matrix patterns in experimental *B. subtilis* biofilms

We wanted to know if *B. subtilis* biofilms did exhibit patterns like those predicted by the model, and if we could validate key features of the model experimentally. To experimentally observe the formation of motile-matrix patterns, we grew *B. subtilis* biofilms in a microfluidic device confined to a thickness of 1-3 cell layers(24). We used a strain with separate transcriptional fluorescent reporters for promoters of motility genes and matrix production genes(10). After roughly 16 hours of growth, biofilms organised into a distinctive pattern much like that predicted by the model, with a fractal-like population of motile cells surrounded by matrix-producing cells (Fig. 6A). The interface pattern remained unchanged for several hours of exterior growth as predicted by the model.

**Figure 6.**
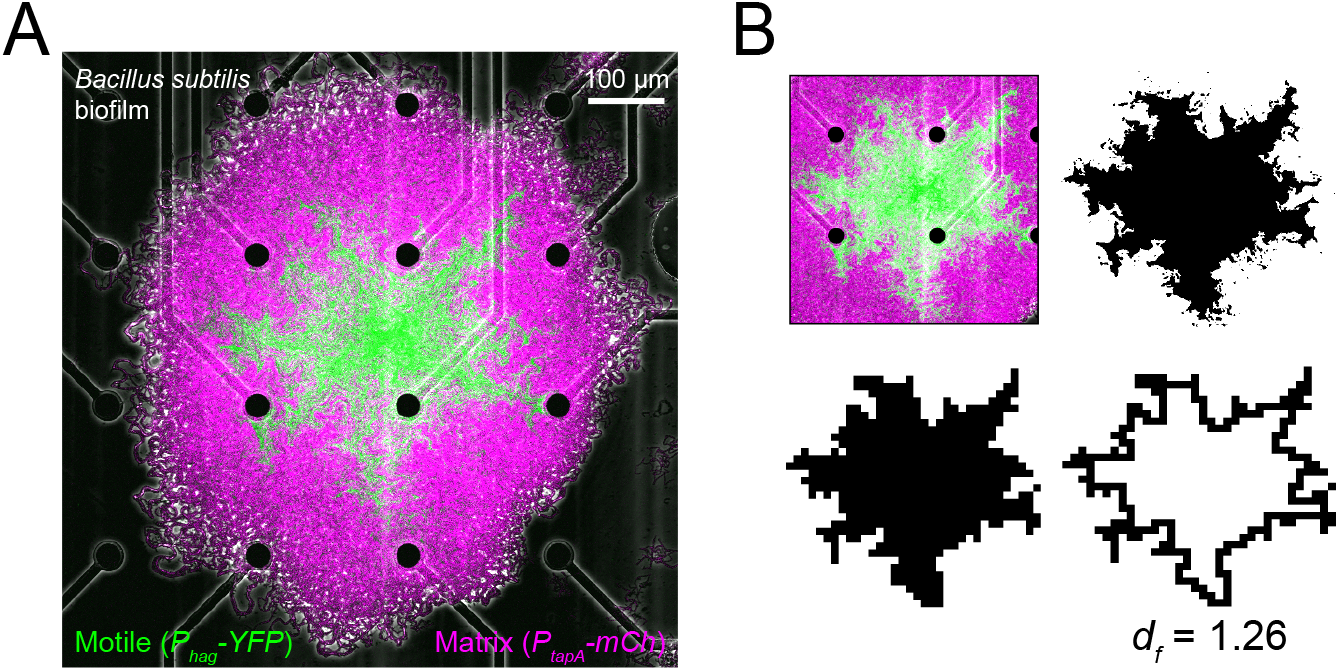
Phenotypic Patterning in a *Bacillus subtilis* Biofilm. (A) Fluorescence image of a *B. subtilis* biofilm grown in a microfluidic device. False colour green is the signal from a reporter on motility genes (*P*_*hag*_ −*YFP*). Magenta is the signal from a matrix production reporter (*P*_*tapA*_ −*mCherry*). (B) Fractal dimension analysis of experimental biofilms. Experimental images segmented according to *P*_*hag*_ −*YFP* intensity, down-sampled to the same resolution as simulations, and fractal dimension is computed with box counting. The fractal dimension is *d*_*f*_ = 1.26 ± 0.03 (*N* = 4)

To determine the experimental fractal dimension, we thresholded biofilm images, found the outline of the interior motile population, and performed box-counting. To account for the fact that a lattice site corresponds to a coarse-grained packet of cells, we down-sampled our experimental images to make them the same resolution as the model (Fig. 6B). We found that the experimental fractal dimension was 1.26 ± 0.03 (*N* = 4), which is within the range predicted by the model (Fig. 3C).

To further compare the experimental properties with the model output, we sought to compute the average radii of the interface and of the biofilm when the pattern freezes, i.e., *r* and *R* of Fig. 5. Because the fluorescent reporters of our experimental strain did not exhibit enough initial fluorescence for segmentation, we could not experimentally measure Δ*r*. However, we were able to estimate the ratio *r/R* for the *N* = 4 biofilms observed, and the results are shown in Fig. 5D (pink stars). We found that the experimental values were cloes together and generally fell within the range of model outputs, tending toward the upper right of the (*d*_*f*_, *r/R*) plot.

These experimental results do not demonstrate that the mechanical mechanisms of our model drive phenotypic pattern formation in *B. subtilis*, but they show that our model predictions are quantitatively consistent with the properties of experimental biofilms.

## DISCUSSION

We have presented a simple, agent-based model that accounts for multiple features of patterns formed by motile and matrix cell phenotypes in *B. subtilis* colonies. Specifically, the model addresses two key questions: 1) how do motile-matrix patterns depend on initial conditions? 2) How do mechanical interactions between heterogeneous cell types contribute to patterning? On the first question, we found that in uncolonised regions of space, our mechanical growth model’s dynamics create patterns of motile cells that are robust to the initial arrangement of cells. This arises from the model spatially distributing cells not only through growth and inheritance, but also by mechanical shoving during cell replication. For three different starting arrangements, we found that the limiting pattern depended strongly on the initial condition within the initial inoculum radius, but beyond that radius the normalised density of motile cells collapsed for all three initial patterns. On the second question, the model predicted that varying the mechanical properties of growth can dramatically alter phenotypic patterns, but that growth and patterning are statistically constrained: biofilms that produce fractal interfaces exhibit more growth of motile cells. It is not clear what, if anything, is the benefit of creating a fractal interface between cells of different phenotypes. Two possibilities are that creating a fractal interface expands biofilms faster due to excessive cell shoving (Fig. S3A) or that a fractal interface minimises the distance between different cell types (Fig. S3B) to make diffusible resource sharing more efficient.

Our model greatly simplifies *B. subtilis* colony growth. Among other things, the model does not take into account nutrient availability, phenotypic switching, cell shape, or the fact that cells grow continuously and not in discrete time. However, despite these simplifications, it makes predictions that are consistent with experimental data and demonstrates a potential mechanical mechanism for spatial distributions of phenotypes within biofilms. Furthermore, our model establishes a starting point for more detailed agent-based simulations that take into account other factors like active cell movement, metabolism, or cell shape.

Our finding that the final pattern only depends on the initial conditions at low radii, and beyond this the resulting pattern is mechanistically determined, is something that we could not have found from experiments. The reason is that there is significant uncertainty in our ability to determine the differentiation state at early time points. This is one advantage to mechanistic modelling: it allows one to validate experiments in particular regimes, and in doing so, makes predictions for regimes not yet accessible.

Attempts to use agent-based or statistical physics models for living systems must take into account an essential feature of cells and organisms: they not only sense their environment, they also change it. This feedback can lead to remarkable phenomena that collectives of cells can take advantage of. Our results capture this phenomenon in a simple way. As biofilm cells grow, they create a constrained environment, altering patterns of growth and leading to a wide variety of potential cell-type patterns.

## MATERIALS AND METHODS

### Model

Code for the model has been posted to the following GitHub repository: https://github.com/caelan-brooks/frac-interface_model

#### Determining the Fractal Dimension of Model Output

The fractal dimension is found using the box counting method. Within the model, first the internal shape is recognised through a recursive shape fill method. From this, the perimeter of this shape is isolated and the box counting method enforced. In order to correctly identify the dimension of the shape, the grids of different box sizes are fit to the perimeter identified. That is, as the biofilm grows, the dimensions of the grids used in analysis will change as well. At each time step, we produce a graph of ln(box size) vs ln(occupied boxes). The negative slope of a regression line is taken as the fractal dimension of the internal motile sub-population. The number of points within this plot is reflective of the number of grids of different box sizes fit to the perimeter. The algorithm attempts to maximise the number of points while limiting the overall dimension of the grid used to cover the internal shape. When repeated over numerous time steps as the biofilm grows and then averaging over hundreds of systems, we are able to produce the results shown in Fig. 3.

### Experiment

We grew biofilms in a microfluidic device as described in (24). Biofilm growth media consisted of liquid MSgg medium containing 5 mM potassium phosphate buffer (pH 7.0), 100 mM MOPS buffer (pH 7.0, adjusted using NaOH), 2 mM MgCl_2_, 700 *µ*M CaCl_2_, 5 *µ*M MnCl_2_, 100 *µ*M FeCl_3_, 1 *µ*M ZnCl_2_, 2 *µ*M thiamine HCl, 0.1 mM sodium citrate, 0.5. % (v/v) glycerol and 0.4% (w/v) monosodium glutamate. Media were made from stock solutions immediately before experiments, and the stock solution of glutamate made fresh daily. We acquired images with a Fluoview FV3000 scanning confocal microscope (Olympus Corp.) using a 10X, 0.3 NA air objective. We performed experiments with a modified *B. subtilis* NCIB3610 strain with Venus yellow fluorescent protein (YFP) fused to the *hag* promoter and mCherry fused to the *tapA* promoter.

### Data Analysis

We analysed images in Fiji(33). In order to accurately compare our model to our experimental data, we first needed to coarse-grain the experimental images–ensuring that the experimental pixel size was comparable to the cell size of our model. To achieve accurate scaling, we first cropped the time-lapse tiff stack for the YFP channel so the saturated fractal filled the entire frame. At this point, select frames were analysed individually and were only chosen after a clear bimodal distribution arose in the YFP channel–the brighter peak representing the high concentration of motile cells inside the motile/matrix interface and the dimmer peak representing regions of the biofilm containing mostly matrix cells. This step ensured that there was minimal bias in thresholding, as images were thresholded simply by selecting the brighter peak of inner motile cells. After filling the holes of the thresholded image, we could resize the entire frame to 40 by 40 pixels, so it was comparable to the size of the saturated fractal in the computer model. Next, we converted the threshold to binary and the fractal was cropped to fit the entire frame–this ensures a more accurate measure of its fractal dimension. We could finally outline the interface and use Fiji’s fractal box count tool, selecting the same sized boxes used for the computer model’s fractal analysis, to identify the fractal dimension of the experimental interface.

## ACKNOWLEDGMENTS

We acknowledge Neil Marya, Patrick Powers, and members of the Larkin Lab for comments on the paper.

## SUPPLEMENTARY MATERIAL

**Supplementary Figure S1.**
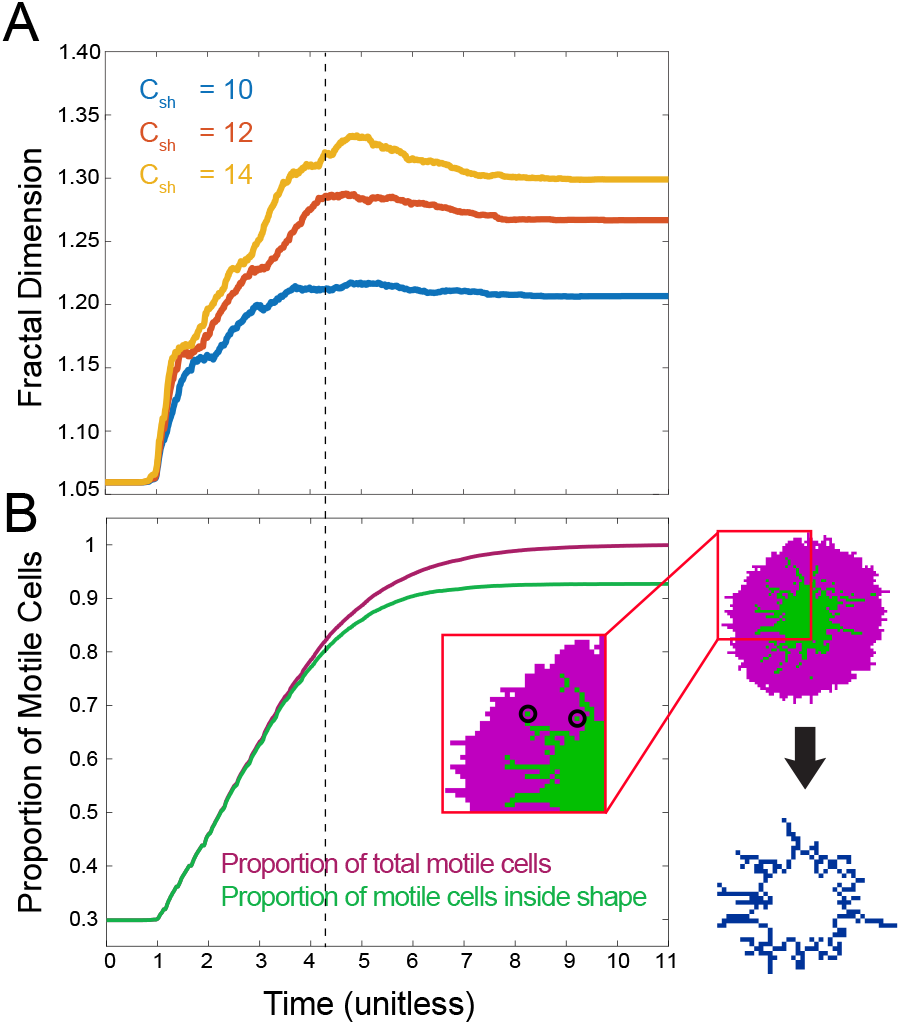
Dynamics of Fractal Dimension. (A) The evolution of the fractal dimension as the simulated biofilm grows (300 simulations averaged). Each line represents biofilm growth under a different shoving capacity. (B) The peak and then decline in fractal dimension is credited to the formation of motile islands. These are motile cells that are not connected to the internal motile shape, but remain stuck inside the matrix producing cells. The split in the two curves signalling the formation of these islands coincides with the peak of the fractal dimension curve (*C*_*sh*_ = 12 used).

**Supplementary Figure S2.**
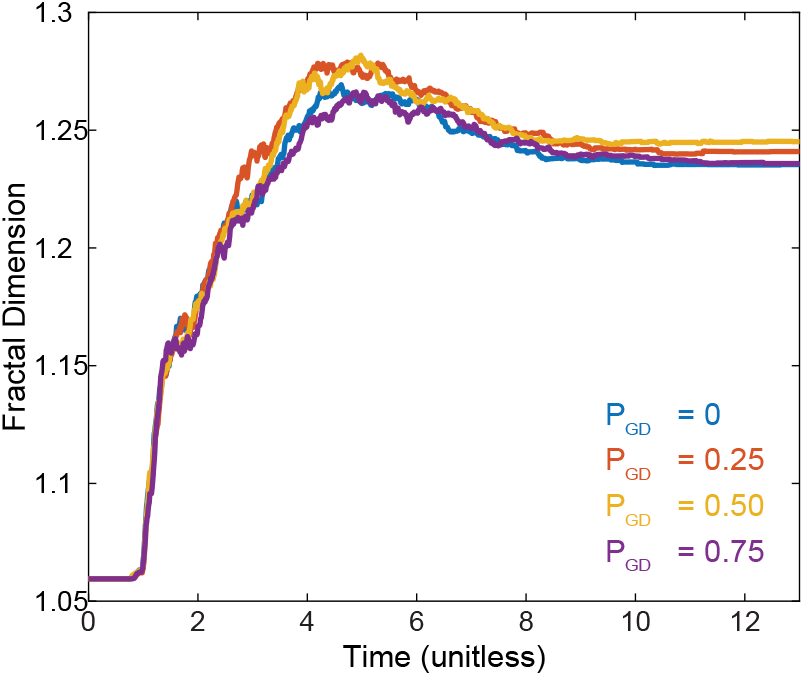
Effects of Preferential Growth Direction. Each line represents a condition in which each cell has a certain probability of growing in the same direction that it previously grew into. The probability for a lineage to continue to choose the same growth direction does not effect how the fractal dimension of the internal motile shape evolves.

**Supplementary Figure S3.**
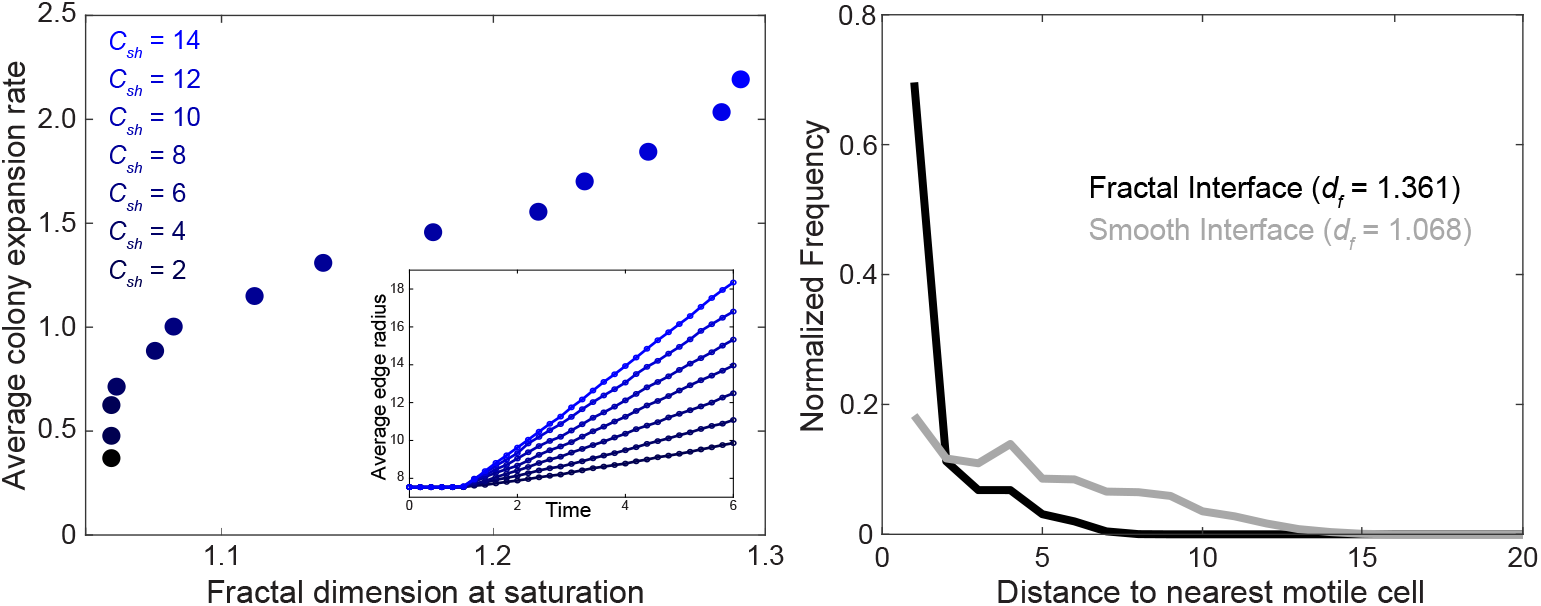
Potential Benefits of a Fractal Interface. (A) Simulating biofilms with a variety of shoving capacities (shades of blue) shows a positive relationship between saturating fractal dimension and overall biofilm growth rate. (B) Biofilms that create a fractal interface (black) have, on average, a much lower distance between cells of different phenotypes.

## REFERENCES

[1] Veening JW, Smits WK, Kuipers OP. Bistability, Epigenetics, and Bet-Hedging in Bacteria. Annual Review of Microbiology. 2008;62(1):193–210. PMID: 18537474. Available from: 10.1146/annurev.micro.62.081307.163002.

[2] Nikolic N, Schreiber F, Dal Co A, Kiviet DJ, Bergmiller T, Littmann S, et al. Cell-to-cell variation and specialization in sugar metabolism in clonal bacterial populations. PLOS Genetics. 2017 12;13(12):1– 24. Available from: 10.1371/journal.pgen.1007122.

[3] Maamar H, Raj A, Dubnau D. Noise in Gene Expression Determines Cell Fate in Bacillus subtilis. Science. 2007;317(5837):526–529. Available from: 10.1126/science.1140818.

[4] Lopez D, Vlamakis H, Kolter R. Generation of multiple cell types in Bacillus subtilis. FEMS Microbiology Reviews. 2008 11;33(1):152–163. Available from: 10.1111/j.1574-6976.2008.00148.x.

[5] Vlamakis H, Chai Y, Beauregard P, Losick R, Kolter R. Sticking together: building a biofilm the Bacillus subtilis way. Nature Reviews Microbiology. 2013 Mar;11(3):157–168. Available from: 10.1038/nrmicro2960.

[6] Kearns DB, Losick R. Cell population heterogeneity during growth of Bacillus subtilis. Genes & Development. 2005;19(24):3083–3094. Available from: http://genesdev.cshlp.org/content/19/24/3083.abstract.

[7] Armbruster CR, Lee CK, Parker-Gilham J, de Anda J, Xia A, Zhao K, et al. Heterogeneity in surface sensing suggests a division of labor in Pseudomonas aeruginosa populations. eLife. 2021 jun;8:e45084. Available from: 10.7554/eLife.45084.

[8] Steinberg N, Keren-Paz A, Hou Q, Doron S, Yanuka-Golub K, Olender T, et al. The extracellular matrix protein TasA is a developmental cue that maintains a motile subpopulation within Bacillus subtilis biofilms. Science Signaling. 2020;13(632). Available from: https://stke.sciencemag.org/content/13/632/eaaw8905.

[9] Stewart PS, Franklin MJ. Physiological heterogeneity in biofilms. Nature Reviews Microbiology. 2008 Mar;6(3):199–210. Available from: 10.1038/nrmicro1838.

[10] Vlamakis H, Aguilar C, Losick R, Kolter R. Control of cell fate by the formation of an architecturally complex bacterial community. Genes & Development. 2008;22(7):945–953. Available from: http://genesdev.cshlp.org/content/22/7/945.abstract.

[11] O’Toole G, Kaplan HB, Kolter R. Biofilm Formation as Microbial Development. Annual Review of Microbiology. 2000;54(1):49–79. PMID: 11018124. Available from: 10.1146/annurev.micro.54.1.49.

[12] Stoodley P, Sauer K, Davies DG, Costerton JW. Biofilms as Complex Differentiated Communities. Annual Review of Microbiology. 2002;56(1):187–209. PMID: 12142477. Available from: 10.1146/annurev.micro.56.012302.160705.

[13] Rosenberg SM. Life, Death, Differentiation, and the Multicellularity of Bacteria. PLOS Genetics. 2009 03;5(3):1–3. Available from: 10.1371/journal.pgen.1000418.

[14] Flemming HC, Wingender J. The biofilm matrix. Nature Reviews Microbiology. 2010 Sep;8(9):623– 633. Available from: 10.1038/nrmicro2415.

[15] Chai Y, Norman T, Kolter R, Losick R. An epigenetic switch governing daughter cell separation in Bacillus subtilis. Genes & Development. 2010;24(8):754–765. Available from: http://genesdev.cshlp.org/content/24/8/754.abstract.

[16] Norman TM, Lord ND, Paulsson J, Losick R. Memory and modularity in cell-fate decision making. Nature. 2013 Nov;503(7477):481–486. Available from: 10.1038/nature12804.

[17] Srinivasan S, Vladescu ID, Koehler SA, Wang X, Mani M, Rubinstein SM. Matrix Production and Sporulation in Bacillus subtilis Biofilms Localize to Propagating Wave Fronts. Biophysical Journal. 2018;114(6):1490–1498. Available from: https://www.sciencedirect.com/science/article/pii/S0006349518301978.

[18] Persat A, Nadell CD, Kim MK, Ingremeau F, Siryaporn A, Drescher K, et al. The Mechanical World of Bacteria. Cell. 2015 May;161(5):988–997. Available from: 10.1016/j.cell.2015.05.005.

[19] Volfson D, Cookson S, Hasty J, Tsimring LS. Biomechanical ordering of dense cell populations. Proceedings of the National Academy of Sciences. 2008;105(40):15346–15351. Available from: https://www.pnas.org/content/105/40/15346.

[20] Smith WPJ, Davit Y, Osborne JM, Kim W, Foster KR, Pitt-Francis JM. Cell morphology drivesspatial patterning in microbial communities. Proceedings of the National Academy of Sciences. 2017;114(3):E280–E286. Available from: https://www.pnas.org/content/114/3/E280.

[21] Rudge TJ, Federici F, Steiner PJ, Kan A, Haseloff J. Cell polarity-driven instability generates selforganized, fractal patterning of cell layers. ACS synthetic biology. 2013;2(12):705–714.

[22] Winkle JJ, Igoshin OA, Bennett MR, Josić K, Ott W. Modeling mechanical interactions in growing populations of rod-shaped bacteria. Physical biology. 2017;14(5):055001.

[23] Narla AV, Borenstein DB, Wingreen NS. A biophysical limit for quorum sensing in biofilms. Proceedings of the National Academy of Sciences. 2021;118(21).

[24] Comerci CJ, Gillman AL, Galera-Laporta L, Gutierrez E, Groisman A, Larkin JW, et al. Localized electrical stimulation triggers cell-type-specific proliferation in biofilms. Cell Systems. 2022. Available from: https://www.sciencedirect.com/science/article/pii/S2405471222001661.

[25] Picioreanu C, Kreft JU, van Loosdrecht MCM. Particle-Based Multidimensional Multispecies Biofilm Model. Applied and Environmental Microbiology. 2004;70(5):3024–3040. Available from: 10.1128/aem.70.5.3024-3040.2004.

[26] Drescher K, Dunkel J, Nadell CD, van Teeffelen S, Grnja I, Wingreen NS, et al. Architectural transitions in ¡i¿Vibrio cholerae¡/i¿ biofilms at single-cell resolution. Proceedings of the National Academy of Sciences. 2016;113(14):E2066–E2072. Available from: 10.1073/pnas.1601702113.

[27] Martínez-Calvo A, Trenado-Yuste C, Lee H, Gore J, Wingreen NS, Datta SS. Interfacial morphodynamics of proliferating microbial communities. bioRxiv. 2023. Available from: 10.23.563665.

[28] Branda SS, Chu F, Kearns DB, Losick R, Kolter R. A major protein component of the Bacillus subtilis biofilm matrix. Molecular Microbiology. 2006;59(4):1229–1238. Available from: 10.1111/j.1365-2958.2005.05020.x.

[29] Stoodley P, Wilson S, Hall-Stoodley L, Boyle JD, Lappin-Scott HM, Costerton JW. Growth and Detachment of Cell Clusters from Mature Mixed-Species Biofilms. Applied and Environmental Microbiology. 2001;67(12):5608–5613.

[30] Vicsek T, Cserző M, Horváth VK. Self-affine growth of bacterial colonies. Physica A: Statistical Mechanics and its Applications. 1990;167(2):315–321. Available from: https://www.sciencedirect.com/science/article/pii/037843719090116A.

[31] Koorehdavoudi H, Paul Bogdan GW, Marculescu R, Zhuang J, Carlsen RW, Sitti M. Multi-fractal characterization of bacterial swimming dynamics: a case study on real and simulated Serratia marcescens. Proceedings of the Royal Society A. 2017;473:20170154.

[32] Gagnepain JJ, Roques-Carmes C. Fractal approach to two-dimensional and three-dimensional surface roughness. Wear. 1986;109(1):119–126. Available from: https://www.sciencedirect.com/science/article/pii/0043164886902577.

[33] Schindelin J, Arganda-Carreras I, Frise E, Kaynig V, Longair M, Pietzsch T, et al. Fiji: an open-source platform for biological-image analysis. Nature Methods. 2012 Jul;9(7):676–682. Available from: 10.1038/nmeth.2019.

